# Aβ promotes amyloidogenic processing of APP through a Go/Gβγ signaling

**DOI:** 10.1101/2021.03.10.434768

**Authors:** Magdalena Antonino, Paula Marmo, Carlos Leandro Freites, Gonzalo Quassollo, Maria Florencia Sanchez, Alfredo Lorenzo, Elena Anahí Bignante

## Abstract

Alzheimer’s disease (AD) is characterized by a cognitive impairment associated to amyloid beta (Aβ) aggregation and deposition in the brain. Aβ is generated by sequential cleavage of the amyloid precursor protein (APP) by β-site APP cleaving enzyme 1 (BACE1) and γ-secretase complex. The mechanisms that underlie exacerbated production of Aβ, favoring its deposition in the brain, is largely unknown. In vitro studies have shown that Aβ aggregates trigger enhanced production of Aβ by a yet non described mechanism. Here, we show that in different cell types, including human neurons derived from induced pluripotent stem cells (iPSC), oligomers and fibrils of Aβ enhance the convergence and interaction of APP and BACE1 in endosomal compartments. We demonstrated a key role of Aβ-APP/Go/Gβγ signaling on the amyloidogenic processing of APP. We show that APP mutants with impaired capacity to bind Aβ or to activate Go protein, are unable to exacerbate APP and BACE1 colocalization in the presence of Aβ. Moreover, pharmacological inhibition of Gβγ subunits signaling with gallein, abrogate Aβ-dependent interaction of APP and BACE1 in endosomes preventing β-processing of APP. Collectively, these findings uncover a feed-forward mechanism of amyloidogenesis that might contribute to Aβ pathology in early stages of AD and suggest that gallein might have clinical relevance.

## INTRODUCTION

Alzheimer’s disease (AD) is the most common neurodegenerative pathology in elderly individuals and is characterized by cerebral deposition of amyloid beta (Aβ). Aβ is a 40-42 amino acid peptide with a natural propensity for self-assembly generating toxic species, including oligomers and fibrils that are difficult to be degraded. An imbalance favoring Aβ production over its clearance leads to accumulation of toxic Aβ species in the brain and the development of the disease^1^. Therefore, identifying molecular mechanisms that enhance Aβ production is key for unveiling the pathophysiology of AD and the adequate targets for rational therapies.

Aβ production begins by shedding the large extracellular portion of amyloid precursor protein (APP) by β-site APP cleaving enzyme 1 (BACE1), the rate-limiting enzyme for Aβ biosynthesis^2^. This cleavage generates a membrane tethered APP C-terminal fragment (β-CFT) and releases a large APP N-terminal soluble fragment (sAPP-β) to the luminal/extracellular space. The intra-membrane cleavage of β-CFT by γ-secretases generates Aβ, liberating the remaining APP intracellular domain (AICD) in the cytosol^3^. Thus, the necessary first step for amyloidogenesis is the convergence of APP and BACE1 in subcellular compartments with the appropriate milieu for BACE1 activity.

In different cell types, amyloidogenic processing of APP has been reported in diverse subcellular compartments, including the Golgi/secretory pathway and endosomes^4–6^. Recent studies revealed that, in neurons, BACE1 processing of APP occurs preferentially in recycling endosomes (RE) in the endocytic pathway^7,8^. However, the signaling process and mechanisms that regulate APP and BACE1 convergence and its consequent β-processing remains poorly understood. Interestingly, previous reports showed that an exogenous application of Aβ assemblies to human leptomeningeal smooth muscle cells or rat neurons in culture enhances amyloidogenic processing of APP and Aβ secretion to the media^9,10^. In line with these findings, local application of pathologic Aβ assemblies can accelerate the spreading of Aβ deposition in the brain of APP transgenic mice^11,12^. These observations suggest that extracellular Aβ assemblies can activate cell surface-signaling mechanisms leading to a feed-forward process thereby enhancing Aβ production and amyloid spreading. Consequently, elucidating the signaling mechanisms underlying this feed-forward process is crucial for understanding the progression of Aβ pathology particularly during early stages of AD.

Heterotrimeric guanine nucleotide-binding proteins (G proteins) are evolutionarily conserved membrane-associated proteins widely used as signal-transduction systems for a myriad of cellular events. G proteins are composed of Gα and Gβγ subunits that directly interact to cell-surface G protein-coupled receptors (GPCR) through Gα subunit. Upon binding to its ligand, the GPCR activates G-protein by exchanging GDP to GTP on Gα subunits, which causes dissociation of Gα from GPCR and from Gβγ subunits. Afterwards, both Gα-GTP and Gβγ subunits can independently activate downstream signaling cascades^13^. Conventional GPCRs are seven transmembrane domain proteins, while unconventional GPCRs are single membrane pass proteins^13^. APP is an unconventional GPCR that can activate Go heterotrimeric protein^14–16^. Significantly, toxic Aβ assemblies require APP to cause neuronal dysfunctions^17–19^. Two different portions of APP ectodomain can interact with Aβ species, the N-terminal portion and the Aβ-juxtamembrane domain^20–23^, both of which can trigger Go activation upon binding to Aβ^24,25^. Surprisingly, no studies have analyzed the implication of Aβ-APP interaction and Go protein signaling for the amyloidogenic processing of APP.

In the present work, we used different cell types, including primary rat hippocampal neurons and human neurons derived from induced pluripotent stem cells (iPSC) and analyzed by advanced microscopy techniques the effect of Aβ assemblies on the convergence and interaction of APP and BACE1 in subcellular compartments. We assessed the role of Aβ-APP-Go signaling on the amyloidogenic processing of APP through the analysis of the effect of the expression of mutant variants of APP, with impaired capacity to bind Aβ or to activate Go protein. Finally, for dissecting the role of Go protein Gβγ subunits in the feed-forward mechanism of amyloidogenesis triggered by Aβ we used gallein, a small molecule that specifically blocks interaction of Gβγ subunits with downstream effectors^26,27^. Our findings reveal that, in human neurons, pathological Aβ assemblies boost β-processing of APP by exacerbating the interaction between APP and BACE1 in RE. This process requires the activation of a signaling pathway that initiates when Aβ interacts with APP, and consequently activates a downstream pathway involving Go protein and Gβγ subunits. Gallein demonstrates robust effectiveness in attenuating the β-processing induced by Aβ. Our work describes a signaling pathway leading to a feed-forward amyloidogenic process with high therapeutic relevance for AD.

## RESULTS

### Aβ enhances APP and BACE1 convergence in recycling endosomes by signaling through Go protein Gβγ subunits in HeLa cells

We have previously characterized in detail that binding of Aβ fibrils (f-Aβ) to cell surface APP specifically activates Go protein^23,24^. To examine the role of APP-Go protein signaling in amyloidogenic processing of APP, we analyzed subcellular localization of fluorescently-tagged APP and BACE1 in culture cells by confocal laser scanning microscopy and performed quantitative colocalization analysis. Cherry fluorescent protein was fused to human BACE1 (BACE1:CH) while yellow fluorescent protein (YFP) was fused to wild type human APP (APP:YFP) or to mutant versions of human APP (Figure 1A). One APP mutant version lacks the extracellular Aβ domain (APPΔβ:YFP), which prevents binding of APP to f-Aβ^21,24,28^. The other APP mutant version contains substitutions of histidines at positions 657 and 658 by glycine and proline (APP_GP_:YFP) respectively, which impairs the ability of APP for activating Go protein signaling^14,29^. We co-transfected HeLa cells with BACE1:CH and APP:YFP, APPΔβ:YFP or APP_GP_:YFP and assessed subcellular localization of APP and BACE1(Figure 1 B-C). We found that APP:YFP distributed along the cytoplasm with a reticular and punctate appearance, which is consistent with the distribution of APP along the secretory and endocytic pathway, localizing in the endoplasmic reticulum, Golgi and endosomes^5,6,30^. We observed that APP_GP_:YFP depicted similar pattern of subcellular localization than wild type APP (Figure 1C). Subcellular distribution of APPΔβ:YFP showed a slightly different appearance with an apparently larger and more defined punctate pattern (Figure 1C), which could reflect the absence of the Aβ sequence that in neurons mediates sorting of APP to axons^31^. However, by quantitative colocalization analysis with adequate compartment markers, we found no significant differences in the distribution of APPΔβ:YFP in recycling endosomes (RE) and Golgi apparatous in basal conditions (Figure 1 and Supplementary Figure 1B,E), underscoring that this mutation did not grossly affect subcellular distribution of the protein in non-polarized HeLa cells. On the other hand, BACE1:CH depicted a punctate expression throughout the cytoplasm, which is consistent with its enrichment in the endosomal compartment^32^. Interestingly, in control condition BACE1:CH distribution was similar regardless of whether it was co-expressed with wild type or mutant variants of APP (Figure 1C,J Supplementary figure 1F).

**Figure 1.**
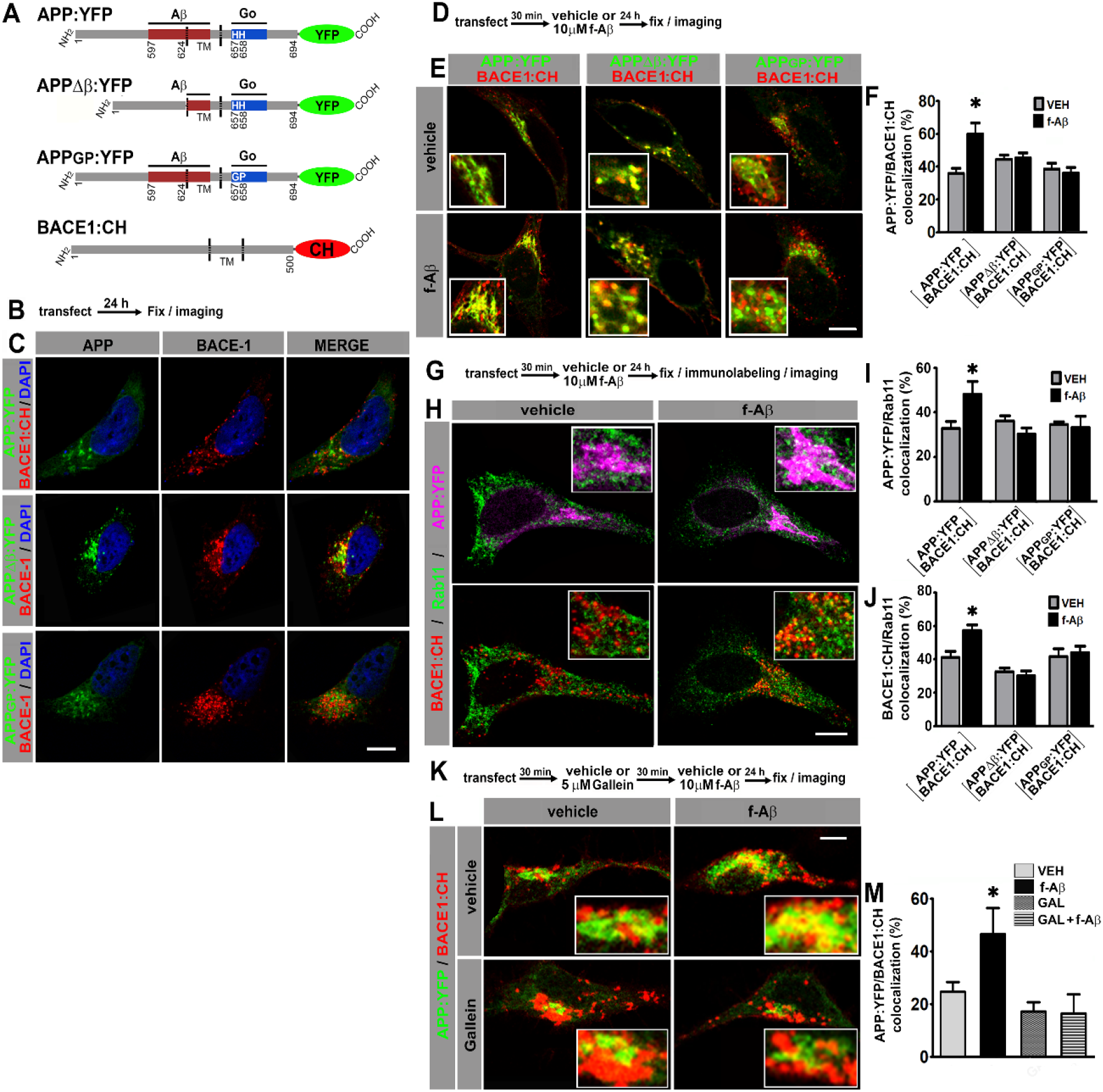
Fibrillar Aβ increases colocalization of APP:YFP and BACE1:CH in recycling endosomes through a Go protein Gβγ subunits-dependent signaling pathway. A. Schematic of variants of APP695 fused to yellow fluorescent protein (YFP) and BACE1 fused to cherry fluorescent protein (CH). Aβ and Go protein-interacting domains are indicated as red and blue boxes, respectively. Amino acid numbering for APP and BACE1 are indicated. TM, transmembrane domain. B. Experimental diagram for C. C. Confocal images of HeLa cells co-expressing the indicated proteins. D. Experimental diagram for E and F. E. Confocal images of HeLa cells co-expressing the indicated proteins. F. Manders coefficients of colocalization of APP:YFP, APPΔβ:YFP or APP_GP_:YFP and BACE1:CH expressed as percent. (n = 56) *: p ≤ 0.001 vs VEH by ANOVA followed by LSD post hoc test. G. Experimental diagram for H-J. H. Confocal images of HeLa cells co-transfected with the indicated proteins and immunolabeled with antibody against Rab11 followed by Alexa Fluor 633-conjugated anti-rabbit antibody. Upper panel shows pseudocolor images for APP:YFP (magenta) and Rab11(green) and lower panel for BACE1:CH (red) and Rab11 (green). I. Manders coefficients of colocalization of APP:YFP and Rab11 expressed as percent. (n = 62 between 8-11 for each condition) *: p ≤ 0.005 vs VEH by ANOVA followed by LSD post hoc test. J. Manders coefficients of colocalization of BACE1:CH and Rab11 expressed as percent.(n = 62; between 8-11 for each condition) * p = 0.001 vs VEH by ANOVA followed by LSD post hoc test. K. Experimental diagram for L and M. L. Confocal images of N2a cells co-expressing the indicated proteins. M. Manders coefficients of colocalization of APP:YFP and BACE1:CH expressed as percent. (n = 29; between 7-8 for each condition) *: p = 0.004 vs GAL + f-Aβ and p = 0.031 vs VEH by ANOVA followed by LSD post hoc test. In all cases data represent mean ± S.E.M. Scale bar 10 μm. VEH: vehicle (control); f-Aβ: fibrillar Aβ1-42; Gal: gallein. Inserts are 2.5 x enlargement of the corresponding image.

Next, we evaluated the effect of f-Aβ on APP and BACE1 colocalization in HeLa cells transfected with BACE1:CH and APP:YFP, APPΔβ:YFP or APP_GP_:YFP (Figure 1D). Notably, we found that f-Aβ significantly augmented colocalization of BACE1:CH with APP:YFP, but did not affected colocalization of BACE1:CH with APPΔβ:YFP or with APP_GP_:YFP (ANOVA F(5,51) = 4.77, p = 0.001; Figure 1E,F). Therefore, in HeLa cells f-Aβ enhances colocalization of wild type APP and BACE1, an effect that requires the Aβ juxtamembrane sequence of APP and the histidine doublet required for activating Go protein signaling.

Although interaction of APP with BACE1 and amyloidogenic processing may occur in different subcellular compartments, we focused on RE and Golgi/secretory pathway. Recent evidence showed that neuronal activity induces β-processing of wild type APP preferentially in RE^7^, while pathogenic Swedish APP mutation (APPSWE) dramatically enhances amyloidogenic processing in the Golgi apparatus/secretory pathway^33,34^. To test whether f-Aβ enhances APP and BACE1 colocalization in RE and/or in the Golgi-secretory pathways, we co-transfected HeLa cells with APP:YFP and BACE1:CH. Afterwards, we treated the cultures with vehicle or f-Aβ, and 24 h later we performed immunolabeling of Rab11 or GM130, which are resident proteins of RE and Golgi apparatus, respectively^35,36^ (Figure 1G). We observed that f-Aβ treatment significantly enhanced simultaneous colocalization of APP:YFP with Rab11 (ANOVA F(5,57)=2.907, p = 0.021; Figure 1H,I) and BACE1:CH with Rab11 (ANOVA f(5;57)=7.795; p < 0.001; Figure 1H,J). On the contrary, we observed that f-Aβ treatment selectively enhanced colocalization of APP:YFP with GM130 (ANOVA f(3,37)=9.303; p < 0.001; Supplementary Figure 1D,E) but did not affect colocalization of BACE1:CH with GM130 (ANOVA F(3,37)=1,8; p = 0.164; Supplementary Figure 1 D,F). Notably, in cells co-transfected with BACE1:CH and APPΔβ:YFP or APP_GP_:YFP we found that f-Aβ treatment was not able to alter colocalization of either APP or BACE1 with Rab11 (Figure 1 I,J and Supplementary Figure 1A-C). Moreover, in cells transfected with APPΔβ:YFP and BACE1:CH colocalization with GM130 was not altered either (Supplementary Figure 1E,F). Thus, in HeLa cells f-Aβ selectively enhances colocalization of APP and BACE1 in Rab11-positive RE, which requires APP-juxtamembrane Aβ domain for binding to f-Aβ and the integrity of APP sequence for activating Go protein signaling.

We have recently found that Gβγ subunits signaling, rather than Gαo, mediates APP-dependent toxicity of f-Aβ in primary neurons^37^. Thus, to determine whether f-Aβ enhances colocalization of APP and BACE1 by activating an APP-dependent signaling through Go protein and Gβγ subunits we used gallein, a small molecule that specifically binds to Gβγ subunits complex preventing its interaction with downstream signaling effectors^26,27^. We applied vehicle, gallein and f-Aβ to N2a cells that were previously co-transfected with APP:YFP and BACE1:CH. Thereafter, we assessed colocalization after 24 h (Figure 1K). We found that gallein completely abrogated enhanced colocalization of APP:YFP and BACE1:CH in cultures treated with f-Aβ (ANOVA F(3,26) = 4.4791; p = 0.012; Figure 1L M). Therefore, in HeLa cells and in N2a cells, f-Aβ enhances colocalization of APP and BACE1 in RE by activating Go protein Gβγ subunits signaling.

### Aβ enhances convergence of endogenous APP and BACE1 in recycling endosomes in primary rat hippocampal neurons by signaling through Gβγ subunits

APP overexpression can generate non-physiological cellular states, leading to artefactual cellular responses and phenotypes^38,39^. To examine if enhanced colocalization of APP and BACE1 in RE stimulated by f-Aβ is physiologically relevant, we analyzed the effect of f-Aβ on APP-BACE1 colocalization in primary hippocampal neurons expressing endogenous protein levels. We treated rat hippocampal cultures of 14 DIV with vehicle or f-Aβ and performed immunofluorescence and quantitative colocalization analysis (Figure 2A). Similar to transfected cells, we found that f-Aβ significantly enhanced colocalization of endogenous APP (t = −2.270; p= 0.038; Figure 2 B-D) and BACE1 (t = −3,093; p = 0.005; Figure 2 E-G) with Rab11. Furthermore, we found that treatment with gallein (Figure 2H) abolished the enhanced colocalization of APP and BACE1 induced by f-Aβ both, in neuronal somas (ANOVA F(1,44)=4.248; p = 0.045; Figure 2 I-K) and neurites (ANOVA F(1,32) = 5.629; p = 0.024; Figure 2 J,L). Collectively, these results demonstrate that in different cell lines and in primary neurons f-Aβ raises the convergence of APP and BACE1 in RE through a APP/Go/Gβγ signaling.

**Figure 2.**
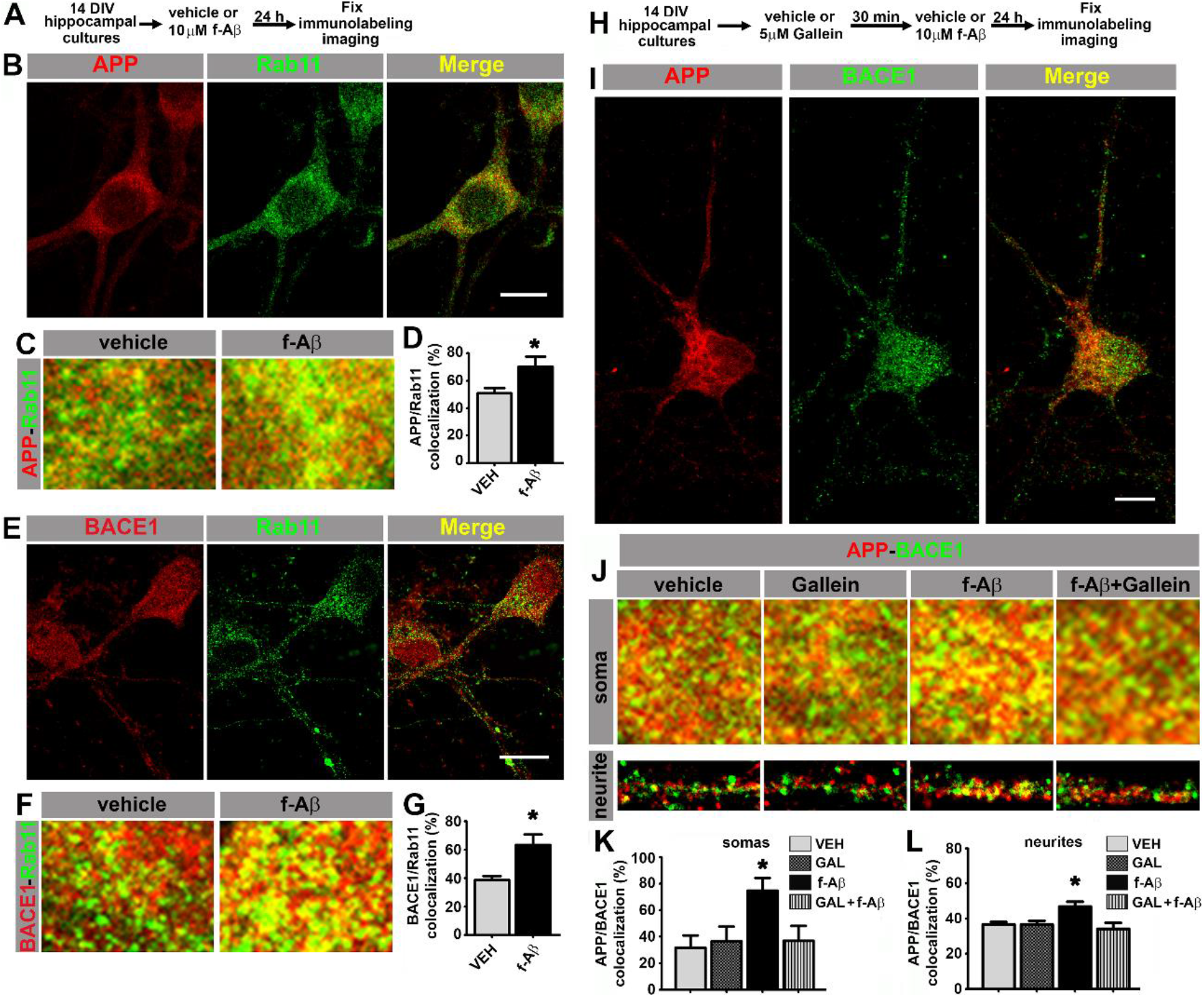
Fibrillar Aβ increases colocalization of endogenous APP and BACE1 in recycling endosomes through Gβγ subunits signaling in hippocampal neurons. A. Experimental diagram for B-G. Rat hippocampal cultures were immunolabeled with anti-APP (clone Y188) and anti-Rab11 (B-D) or anti-BACE1 (clone 3C1C3) and anti-Rab11 (E-G). B and E. Representative confocal images of control cultures. C and F. Enlargements (3 x) of neuronal cell soma of cultures treated with the indicated conditions. D. Manders coefficients of colocalization of APP and Rab11 expressed as percent. (n = 17; veh = 8, f-Aβ = 9) *: p = 0.038 by t test. G. Manders coefficients of colocalization of BACE1 and Rab11 expressed as percent. (n = 25; veh = 12, f-Aβ = 13) *: p = 0.005 by t test. H. Experimental diagram for I-L. Rat hippocampal cultures were immunolabeled with anti-APP (clone Y188) and anti-BACE1 (clone 3C1C3). I. Representative confocal images of control immunolabeled cultures. J. Enlargements (3 x) of neuronal cell soma and neurites of cultures with the indicated conditions. K. Manders coefficients of colocalization of APP and BACE1 in somas, expressed as percent. (n = 48; 12 for each condition) *: p < 0.05 vs all other conditions by ANOVA followed by Fisher LSD post hoc test. L. Manders coefficients of colocalization of APP and BACE1 in neurites, expressed as percent. (n = 36; 8-10 for each condition) *: p < 0.05 vs all other conditions by ANOVA followed by Fisher LSD post hoc test. In all cases data represent mean ± S.E.M. Scale bar 10 μm. VEH: vehicle (control); f-Aβ: fibrillar Aβ1-40; Gal: gallein.

### Aβ increases direct interaction of APP and BACE1 and amyloidogenic processing by signaling through Gβγ subunits

Amyloidogenic processing requires physical interaction of APP with BACE1. To determine whether colocalization induced by f-Aβ is effectively associated with increased physical interaction between APP and BACE1, we used fluorescence bimolecular complementation (BiFC). BiFC is a validated technique for detecting interaction between a protein pair, in which one protein of the pair is fused to the N terminal fragment of the Venus fluorescent protein (VN) and the other protein is fused to the complementary C terminus (VC). VN or VC are not fluorescent, however when the protein pair physically interacts, Venus is reconstituted and becomes fluorescent^40^. An additional advantage of this technique is that complementation of Venus is irreversible and therefore transient protein-protein interactions are stabilized, allowing its visualization. Here, we used expression vectors for BiFC designed and extensively characterized for studying APP and BACE1 interaction in living neurons^8^. Briefly, one vector drives the expression of wild type APP tagged with VN (APP:VN) while the other induces expression of BACE1 tagged with VC (BACE1:VC) (Figure 3A). In order to determine the effect of f-Aβ and the role of APP-Go protein Gβγ subunits signaling on the physical interaction between APP and BACE1, we co-transfected N2a cells with APP:VN and BACE1:VC vectors. Thereafter, we treated the cultures with corresponding vehicles, gallein and f-Aβ and we assessed BiFC intensity after 24 h (Figure 3B). We noticed that in control conditions APP-BACE1 BiFC depicts discrete vesicular fluorescent dots throughout the cytoplasm. Notably, f-Aβ treatment significantly enhanced BiFC, and gallein completely restored BiFC to control levels (Figure 3C). Quantitative assessment confirmed a significant enhancement of APP-BACE1 BiFC intensity in f-Aβ-treated cultures and its reversion by gallein (ANOVA F(2,46)= 5.058; p= 0.001; Figure 3D).

**Figure 3.**
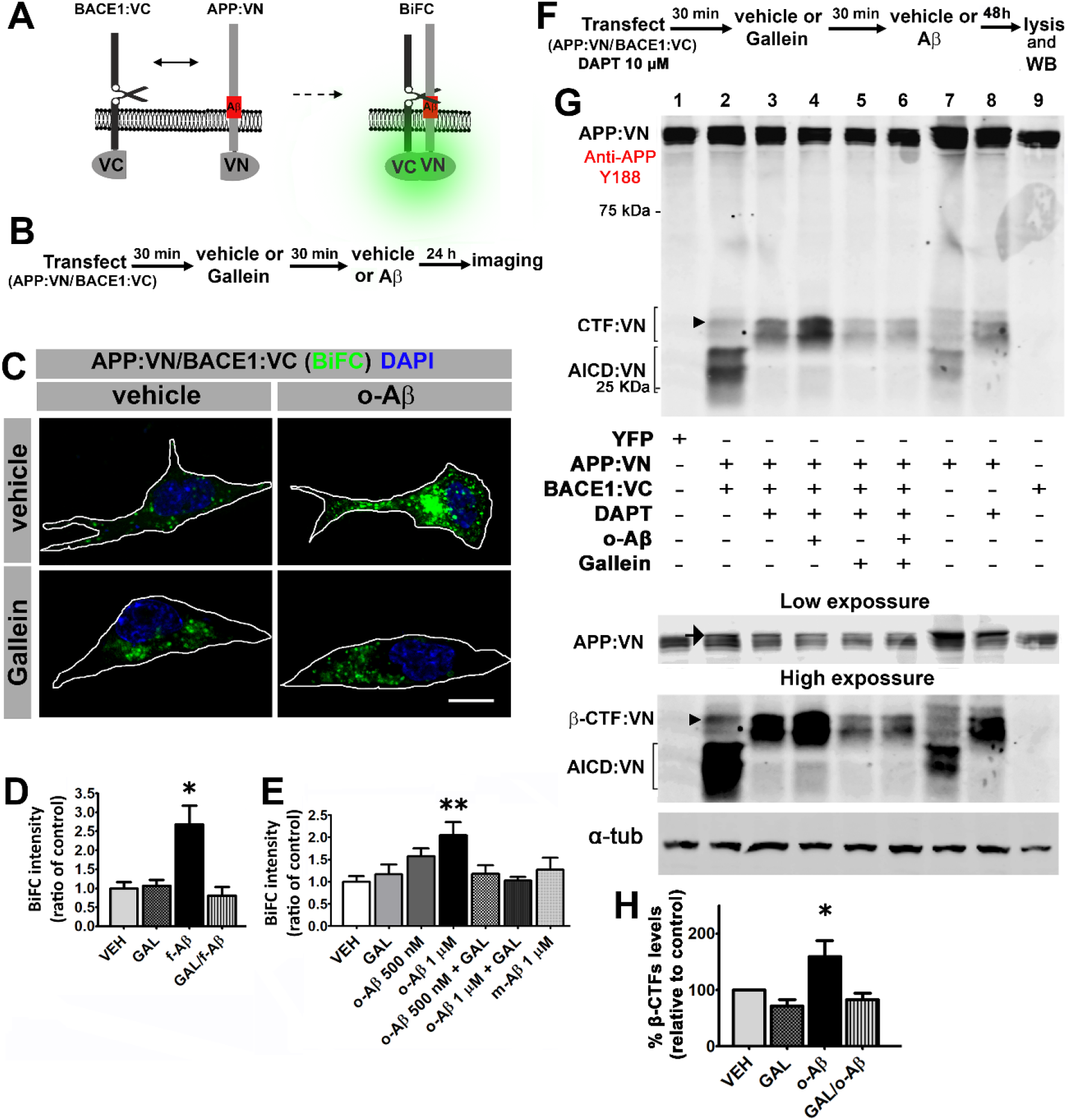
Aβ assemblies enhance physical interaction of APP and BACE1 and amyloidogenic processing through a Gβγ dependent signaling pathway. A. Schematic of bimolecular fluorescent complementation (BiFC) assay for detecting APP-BACE1 interaction. Interaction of APP and BACE1 fused to complementary fragments (VN and VC) of Venus fluorescent protein reconstitutes fluorescence. B. Experimental diagram for C-E. C. Representative fluorescent images of BiFC in N2a cultures co-transfected with plasmid encoding to APP:VN and BACE1:VC and treated with the indicated conditions. D. BiFC intensity expressed as ratio of control. (n = 52; 9-20 for each condition) *: p < 0.001 vs all other conditions by ANOVA followed by Fisher LSD post hoc test. E. BiFC intensity expressed as ratio of control. (n = 109; 12-18 for each condition) *: p < 0.01 vs VEH and vs o-Aβ 1uM+ GAL by ANOVA followed by Fisher LSD post hoc test. Scale bar 10 μm. F. Experimental diagram for G and H. G. Cell lysates of N2a cells co-transfected with APP:VN and BACE1:VC with the indicated conditions were analyzed by western blot analysis with antibody Y188 that recognize the C-terminal domain of APP. Bands corresponding to CTF:VN and AICD:VN are indicated by brackets. Molecular weight markers are indicated in kDa. Low and high exposure are shown down. Arrow in the low exposure panel point to full length APP:VN. Arrowhead point to β-CTF:VN. Loading control α-tubulin is shown in the bottom. H. Quantitative densitometry analysis of β-CTF:VN bands after normalization of their levels to those of corresponding loading control band. (n = 4) *: p < 0.05 vs all other conditions by ANOVA followed Fisher LSD post hoc test. In all cases values denote Means ± S.E.M

Conversion of physiologic monomeric Aβ to pathologic Aβ assemblies, including Aβ oligomers and fibrils, plays a key role in neuronal dysfunction in AD. Therefore, it is important to determine whether non-toxic and toxic Aβ species can enhance APP-BACE1 interaction. We generated non-aggregated soluble Aβ (m-Aβ) and aggregated soluble oligomers (o-Aβ)^41^ and characterized these Aβ species by dot blot with 4G8, NU1 and NU4 antibodies (Supplementary Figure 3). N2a cultures transfected with APP:VN and BACE1:VC were treated with m-Aβ or pathologic o-Aβ. APP-BACE1 BiFC intensity was analyzed after 48 h. Notably, we found that m-Aβ did not altered BiFC (Figure 3E). However, similar to f-Aβ, o-Aβ induced a dose dependent increase in BiFC that reached significance at 1 μM. Moreover, gallein efficiently abrogated the enhancement in BiFC induced by o-Aβ (ANOVA F(3,106) = 3.663; p = 0.016; Figure 3 E). Thus, pathologic assemblies of Aβ, including oligomers and fibrils, enhance physical interaction of APP and BACE1 by a mechanism dependent Gβγ subunits signaling.

To find out whether APP-BACE1 interaction genuinely correlates with β-processing of APP, we expressed APP:VN and BACE1:VC and analyzed BACE1-cleaved C-terminal fragment of APP (β-CTF) by western blot. If BACE1:VC effectively cleaves APP:VN protein, a β-CTF linked to the 17–19 kDa VN protein (β-CTF:VN) should be generated. In addition, cleavage of β-CTF:VN by γ-secretase should generate APP:VN intracellular domain fragments (AICD:VN). Hence, if γ-secretase activity is blocked by the inhibitor DAPT (10 μM), β-CTF:VN should accumulate. We transfected N2a cultures with APP:VN and BACE1:VC. Thereafter, we treated cultures with corresponding vehicles, DAPT, gallein, o-Aβ. After 48 h we harvested the cells and analyzed the lysates by western blot with antibody against APP C-terminal domain (Figure 3F). We observed that in control condition, expression of APP:VN alone produced two groups of bands (Figure 3G lane 7). The lower group was abolished by DAPT (Figure 3G lane 8), indicating that it corresponds to AICD:VN. On the contrary, the upper group was enhanced by DAPT-treatment indicating that it corresponds to CTF:VN. In these experimental conditions, β-processing is due to endogenous BACE1. Co-expression of BACE1:VC with APP:VN (Figure 3G line 2), increased the intensity of both groups of bands, consistent with an enhanced β-processing of APP:VN by BACE1:VC. Again, DAPT completely abrogated γ-secretase processing preventing generation of AICD:VN and leading to enhanced accumulation of CTF:VN (Figure 3G lane 3). These bands undoubtedly are BACE1:VC-dependent and therefore correspond to β-CTF:VN (Figure 3G arrowhead). Importantly, treatment with o-Aβ dramatically increased β-CTF:VN (Figure 3G line 4) which was abolished by gallein (Figure 3G line 6) (F(1,14)= 5.501 p= 0.015; Figure 3H). Finally, gallein *per se* also appears to reduce basal generation of CTF:VN, but this effect lacks statistical significance (Figure 3G line 5 vs line 3). Collectively, these findings demonstrate that pathologic assemblies of Aβ activates Go protein Gβγ subunits signaling promoting the physical interaction of APP and BACE1 leading to enhanced amyloidogenic processing, which might contribute to uphold amyloid pathology in AD.

### Aβ promotes an increase in the interaction between APP:VN and BACE-VC in recycling endosomes of human neurons by signaling through Gβγ subunits

For further appraising the pathophysiological significance of our observations and the clinical potential of gallein for AD, we analyzed the effect of o-Aβ and gallein on the physical interaction between APP and BACE1 in human neurons. Neuronal cultures derived from human iPSC were generated (Supplementary Figure 3) and transfected at 10 DIV with expression vectors for APP:VN and BACE1:VC. Thereafter we incubated the cultures with vehicle, gallein or o-Aβ. After 48 h of treatment cultures were fixed and immunostained against MAP2 and Rab11 proteins (Figure 4A-B). Quantitative assessment of BiFC intensity showed that o-Aβ induced a significant increase in APP:VN and BACE1:VC interaction as evidenced by enhanced BiFC intensity in both, the cell soma (ANOVA F(1,23)=4.429 p=0,046) and dendrites (ANOVA F(1,48)=5.213, p=0,027), an effect that was abrogated by gallein (Figure 4B-E). Notably, o-Aβ treatment significantly augmented APP-BACE1 BiFC and its colocalization in Rab11-positive recycling endosomes, which again was abolished by gallein (ANOVA F(1,23) = 4.63, p = 0.042; Figure 4F,G). Therefore, in human neurons o-Aβ enhances physical interaction of APP and BACE1 in RE by a mechanism that involves Gβγ signaling. Collectively, these findings uncover the signaling mechanism through which Aβ triggers a feed-forward process of amyloidogenesis that might contribute to an imbalance between amyloid production and clearance in early stages of Aβ pathology, and suggest that gallein might have clinical relevance for AD.

**Figure 4.**
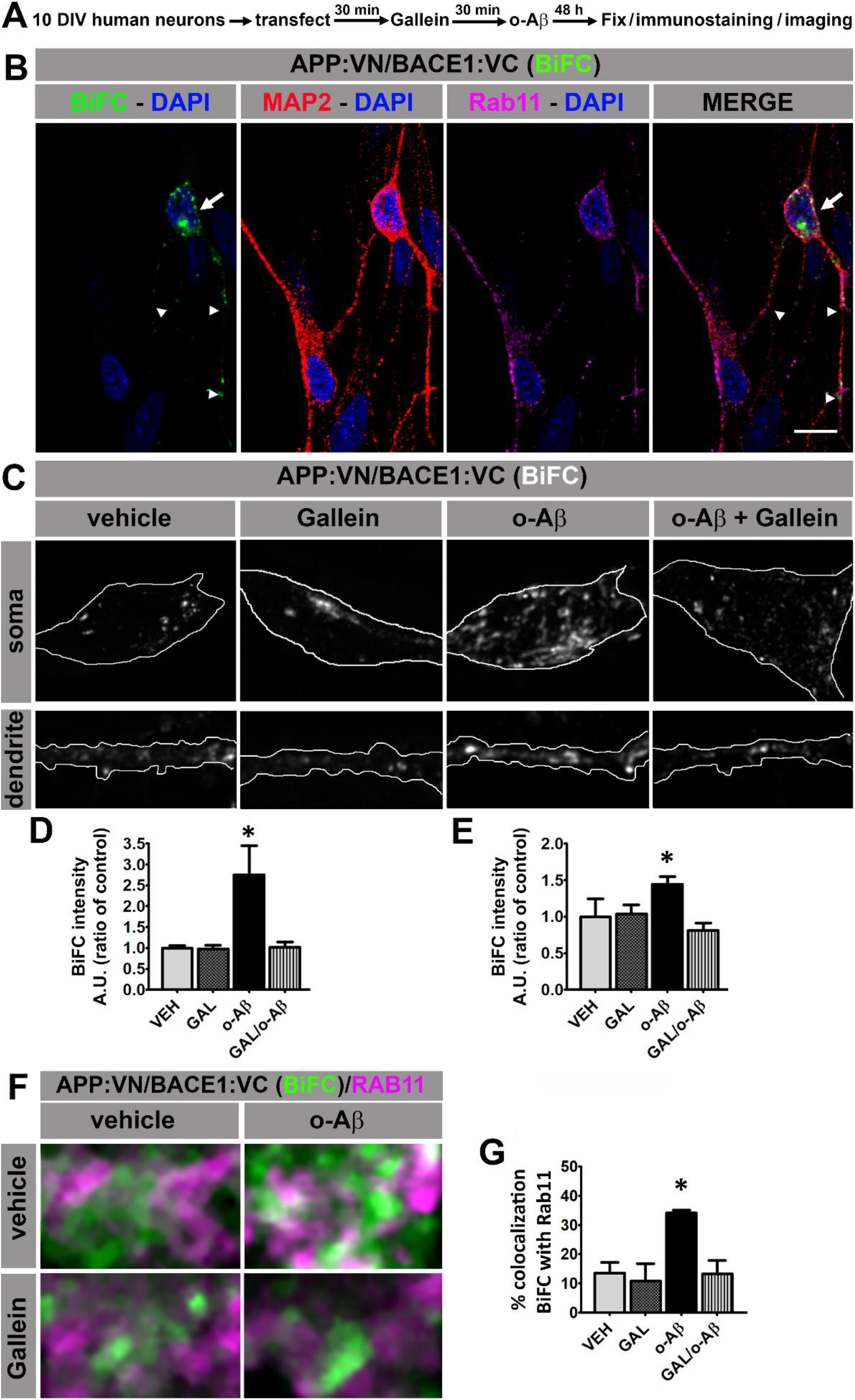
Aβ oligomers promote interaction of APP:VN and BACE1:VC in recycling endosomes of human neurons through a Gβγ subunits dependent signaling pathway. A. Experimental diagram for B-G. B. Confocal image of 12 DIV human neurons immunolabeled with antibodies to MAP2 and Rab11 depicting BiFC of APP:VN and BACE1:VC. BiFC in cell body and dendrites is indicated with arrows and arrowheads, respectively. Nuclei were stained with DAPI. C. Representative images of somas and dendrites of 12 DIV human neurons treated with the indicated conditions showing BiFC of APP:VN and BACE1:VC. D. Intensity of BiFC of APP:VN and BACE1:VC in somas expressed as ratio of control. (n = 27; 6-9 for each condition)*: p < 0.001 vs all other conditions by ANOVA followed Fisher LSD post hoc test. E. Intensity of BiFC of APP:VN and BACE1:VC in dendrites expressed as ratio of control. (n = 52; 11-14 for each condition) *: p < 0.05 vs all other conditions by ANOVA followed Fisher LSD post hoc test. F. Representative images of magnified area of somas of human neurons treated with the indicated conditions and depicting BiFC of APP:VN and BACE1:VC (green) and endogenous Rab11 (magenta). White color indicates colocalization of BiFC and Rab11. G. Manders coefficient of colocalization of BiFC and Rab11 expressed as percentage. Data represent Means ± S.E.M. (n = 27; 6-9 for each condition). *: p < 0.01 by ANOVA followed by Fisher post hoc test. Scale bar 10 μm.

## DISCUSSION

This work elucidates the cellular and molecular mechanism through which pathological Aβ assemblies stimulate the first and most critical step for Aβ production: the encounter and processing of APP by BACE1 in endosomal compartments for the generation of βCFTs. We demonstrate that in diverse cellular types, including human neurons, pathologic Aβ assemblies drive an increased APP/BACE1 interaction and amyloidogenic processing by activating an APP-dependent signaling pathway engaging Gβγ subunits of Go protein. Gallein, a specific Gβγ inhibitor, greatly attenuates the exacerbated β-processing of APP triggered by Aβ, demonstrating the potential value of our findings for AD therapeutics.

AD pathology develops with a lengthy and silent preclinical phase in which Aβ deposition disseminates throughout the brain in a systematic and stereotyped fashion^42,43^. The spreading of Aβ in the human brain is reminiscent to a prion mechanism of propagation. Thus, in mice models of AD, local application of a small seed of pathologic Aβ assemblies accelerates Aβ plaque formation throughout the brain^11^, which presumably involves transynaptic dissemination of Aβ pathology^44,45^.

Likewise, Aβ aggregates triggers an enhanced Aβ production in neurons^10^, suggesting a feed-forward process contributing to perpetuate Aβ deposition in AD. In this work, we found that Aβ assemblies drive intracellular redistribution and enhanced interaction of APP and BACE1 in RE, increasing amyloidogenic processing. This process requires the integrity of APP as a GPCR that is activated by Aβ assemblies. Experimental evidence indicate that physiological and pathological effects of Aβ requires binding to APP ectodomain^17–25,46^. Here we show that APPΔβ-YFP, which lacks the juxtamembrane Aβ sequence for Aβ binding, was ineffective in promoting convergence of APP and BACE1 in the presence of Aβ assemblies. Similarly, APPGP-YFP, an APP mutant ineffective for activating Go protein^14,29^, completely abrogated APP and BACE1 convergence induced by Aβ. Strikingly, the fact that these APP mutants prevent not only their own subcellular redistribution in response to Aβ, but also that of BACE1, reinforces the role of APP as an unconventional GPCR that responds to Aβ through Go protein-dependent signaling cascade. Furthermore, pharmacological inhibition of Gβγ subunits with gallein avoided the redistribution of APP and BACE1 as well as the generation of βCFTs, strengthening the importance of Gβγ subunits signaling on this process.

Spatial intracellular distribution and traffic of APP and secretases have a pivotal role in amyloidogenic processing and pathogenesis of AD. It was postulated that healthy neurons maintain a fine regulation of intracellular traffic to limit APP and BACE1 proximity and Aβ production^47^. In neurons, APP and BACE1 colocalize in several intracellular compartments of the secretory pathway including the Golgi apparatus. However, β-cleavage of wild type APP might not preferentially occur in the secretory pathway due to the abundance of other substrates with much higher affinity for BACE1^48^. Consistent with this, studies conducted in neuronal cells have found that BACE1 processing of APP occurs mainly in the endocytic pathway and that neuronal activity triggers the convergence of APP and BACE1 in RE of hippocampal neurons, increasing β-cleavage of APP^7,8^. Remarkably, enhanced encounter of APP and BACE1 in RE was also reported in neurons of AD brain, suggesting that pathologic conditions in AD stimulate amyloidogenic processing of APP^7^. Our results provide a mechanistic explanation to the later observation showing that pathological assemblies of Aβ, including oligomers and fibrils, favor colocalization of APP and BACE1 and amyloidogenic processing in RE by activating an APP-dependent Go protein signaling pathway. This feedforward process might contribute to uphold Aβ deposition in AD brain.

We also found that in HeLa cells f-Aβ provoked an increase of APP:YFP in the Golgi apparatus. Although, this was not accompanied by a concomitant increase of BACE1:CH, it cannot be excluded that an increased β-processing of APP could also take place in Golgi/secretory pathway. In fact, β-processing of pathologic Swedish APP mutation, which has higher affinity for BACE1 than wild type APP and causes early onset AD^49^, occurs in the Golgi apparatus and secretory pathway^33,34^. Therefore, it is possible that Aβ assemblies could enhance β-processing of wild type APP in Golgi/secretory pathway by increasing the availability of APP for BACE1 processing in these subcellular compartments. Although further experiments will be required to confirm this possibility, our observations emphasize that Aβ assemblies favor the conditions for amyloidogenic processing of APP.

How does APP/Go/Gβγ signaling triggered by Aβ enhance the interaction of APP and BACE1 in RE? It was reported that just a very low proportion of endocyted APP converges with BACE1 in endosomes for β-processing, avoiding the default route for lysosomal degradation^8,50^. In fact, at physiologically state, ubiquitination of cell surface APP leads to its sorting to intraluminal vesicles (IVLs) for lysosomal degradation through a pathway that involve phosphatidyl inositol-3-phosphate (PI3P) and endosomal sorting subunitss required for transport (ESCRT)^51^. Inhibition of this pathway by targeting the lipid kinase Vps34 leads to accumulation of APP in endosomes and enhanced Aβ production. These authors also reported decreased levels of PIP3 in brains of AD patients and rodent models of AD suggesting a potential relation of this signaling with amyloidogenic processing of APP in the pathology^51^. Modulation of this pathway could also contribute to the enrichment of APP in RE after Aβ treatment. In fact, some members of phosphoinositide 3-kinase (PI3K) family have been identified as targets of GPCR and Gβγ subunits^52–54^. Reduced APP degradation in lysosomes would also explain why Aβ deposition increases cellular APP levels through post-translational mechanisms^9,28^.

Interestingly, pathological Aβ aggregates also increase BACE1 levels in neuronal cultures by a mechanism independent of protein synthesis^55^, which also depends on Aβ binding to APP. In fact, it was found that crosslinking surface APP with monoclonal antibody 22C11, which mimics the interaction of Aβ to APP; enhances Aβ production by inactivating the adaptor protein Golgi-localized gamma ear-containing ARF-binding 3 (GGA3) preventing the sorting of BACE1 to lysosomes for degradation^56^. It is important to mention that, similar to toxic Aβ assemblies, multimerization of cell surface APP by incubation with antibody 22C11 also causes toxicity by activating Go protein signaling^57,58^. Altogether, these observations strengthen our observation and allow us to suggest that binding of Aβ assemblies to APP enhances its amyloidogenic processing through Go protein Gβγ subunits by modulating downstream effectors such as PI3K and GGA3 that redirect endosomal trafficking of APP and BACE1 to endosomes. Regardless of which these effectors are, the evidence presented here demonstrates that an efficient strategy to stop exacerbated Aβ production in AD could be achieved by blocking the Go/Gβγ pathway, and that gallein has therapeutic potential in this regard.

Collectively, our results uncover the signaling that triggers a feed-forward mechanism elicited by Aβ assemblies, exacerbating amyloidogenic processing of APP. Over time, this pathological mechanism would favor Aβ deposition and spreading in AD brain. Therefore, this process might have particular relevance for early stages of both, early and late onset AD (EOAD and LOAD respectively). Hence, targeting the signaling APP/Go/Gβγ pathway may have therapeutic potential for halting Aβ pathology in both forms of the disease. Moreover, based in the evidence that targeting GPCR or their downstream signaling cascade is the most successful pharmacological strategy for dealing with most of the human pathologies^59^, we propose that the intervention of this pathway at different levels could have therapeutic relevance. Effectiveness of gallein in modulating this process in vitro is the first proof of principle that targeting this signaling cascade could be a new therapeutic approach for AD treatment.

## MATERIAL AND METHODS

### List of antibodies and reagents

Antibodies used were the following: rabbit anti-APP C-terminal clone Y188 (1:250; Abcam, Cambridge, UK), mouse anti-BACE1 clone 3C1C3 (1:100; ThermoFisher, Waltham, MA, U.S.); mouse anti-neuron-specific class III β tubulin clone #TuJ1 (1:2000; Abcam, Cambridge, UK), mouse anti-MAP2 clone AP-20 (1:500; Merck Millipore; Burlington, MA, U.S.), rabbit anti-Rab11 (1:50; ThermoFisher, Waltham, MA, U.S.), mouse anti-GM130 clone 35/GM130 (1:500; BD Bioscience, Franklin Lakes, NJ, U.S.), mouse anti-Aβ17-24 clone 4G8 (1:1000; Biolegend, San Diego, CA, U.S.), mouse anti-Aβ oligomeric clones NU1 and NU4 (1:1000; generous gift from Dr Charles G. Glabe, University of California, Irvine, CA, U.S.), mouse anti-Gαo clone A2 (1:500; Santa Cruz Biotechnology, Dallas, TX, U.S.), mouse anti-GFP clone 3E6 (1:1000; Molecular Probes, Eugene, OR, U.S.), mouse α-tubulin clone B-5-1-2 (1:1000; Sigma Aldrich, San Luis, MO, U.S.). For immunofluorescence primary antibodies were labeled with Alexa-conjugated secondary antibodies Alexa 488, 546, 598 or 633 (1:500; Invitrogen, Carlsbad, CA, U.S.), and for western blotting primary antibodies were conjugated with secondary antibodies IR800CW and IR680RD (1:5000, LI-COR, Lincoln, Nebraska, U.S.) and analyzed in IR Oddissey System.

The following reagents were used: Polyethyleneimine 87K (PEI, produced by Dr. Juan M Lázaro-Martinez, Universidad Nacional de Buenos Aires, Argentina), Lipofectamine 2000 (ThermoFisher, Waltham, MA, U.S.) was used according to manufacturer instructions; gallein (Santa Cruz Biotechnology, Dallas, TX, U.S.) and DAPT (Santa Cruz Biotechnology, Dallas, TX, U.S.) were dissolved in DMSO and applied directly to the cultures at the indicated concentrations.

### Preparation of Aβ assemblies

Synthetic Aβ1-42 and Aβ1-40 were obtained from Biopeptide Inc (San Diego, CA, U.S.). Fibrillar Aβ was prepared as previously described (Heredia et al., 2004). Briefly, 1mM Aβ solution was prepared adding 1,1,1,3,3,3-Hexafluoro-2-Propanol (HFIP, Sigma Aldrich, San Luis, MO, U.S) to the vial containing lyophilized powder. Solution was incubated at room temperature (RT) for at least 30 min and aliquoted. Then, tubes were opened and HFIP evaporated at RT allowing a film formation. Films were stored at 20°C. Aβ film was re-suspended in sterile water and 2x phosphate buffered saline (PBS) in consecutive days and incubated at 37°C for 24 h until a concentration of 500 μM. Whole solution was used for treatments. Oligomeric Aβ was prepared as described by Stine et al.(2003). Briefly, Aβ films were obtained as explained above. Then, Aβ stock solution was prepared by adding dimethyl sulfoxide (DMSO, Sigma Aldrich, San Luis, MO, U.S) to the tube until reach a concentration of 5 mM. Solution was vortexed and sonicated for 10 min. Cold phenol-free F-12 cell culture media was aggregated to the tube, diluting to a final concentration of 100 μM Aβ. Monomeric Aβ was prepared by dissolution of Aβ film in sterile water reaching a final concentration of 100 μM Aβ. Aβ1-42 fribrils were used for colocalization studies in cell lines. Hippocampal neurons, which are especially susceptible to toxicity of Aβ1-42 fibrils and oligomers, were treated with the less aggressive Aβ1-40 fibrils. Human neurons were treated with Aβ1-42 oligomers or fibrils.

### DNA constructs

Full-length human APP695 wild-type and mutant forms (APP:YFP, APPΔβ:YFP, APPGP:YFP) were inserted into pEYFP-N3 encoding the yellow fluorescent protein (YFP, Clontech, Mountain View, CA, U.S.). APPΔβ was generated by deletion of the juxtamembrane domain (aa 597-624) of human APP695 and was previously characterized (Sola Vigo et al 2009, Kedikian et al 2010). APPGP contains a double substitution of H657 and H658 by G and P, respectively. The mutations were introduced into human APP695 using the QuickChange Site-Directed Mutagenesis Kit (Stratagene, San Diego, CA, U.S.) and confirmed by sequencing. This double mutation prevents APP-dependent activation of Go protein (Nishimoto et al., 1993; Brouillet et al., 1999). BACE:CH, APP:VN and BACE1:VC were generously provided by Dr. Subhojit Roy, University of California, San Diego, La Jolla, CA, U.S.

### Culture of cell line and rat primary hippocampal neurons

N2a and HeLa cells were cultured in DMEM supplemented with 10% fetal bovine serum (FBS) (Gibco, Thermo Fisher, Waltham, MA, U.S) and Glutamax (Gibco, Thermo Fisher, Waltham, MA, U.S). For primary neuronal hippocampal cultures, pregnant Wistar rats were provided by the vivarium of the Instituto de Investigación Médica Mercedes y Martín Ferreyra (INIMEC-CONICET-UNC). All procedures regarding the use of animals were conducted in accordance with the National Institute of Health Guide for the Care and Use of Laboratory Animals as approved by the Animal Care and Use Committee of the INIMEC-CONICT-UNC. Hippocampal cultures were established as described previously (Heredia et al., 2004). Briefly, the hippocampi of embryonic day 18 fetuses were dissected. After a brief incubation in trypsin, cells were mechanically dissociated with a Pasteur pipette and plated at 100 cells/mm^2^ on poly-L-lysine (0.25 mg/ml, Sigma-Aldrich) coated coverslips in DMEM (Gibco, Thermo Fisher, Waltham, MA, U.S) supplemented with 10% horse serum (Gibco, Thermo Fisher, Waltham, MA, U.S). After 2 h, medium was replaced with Neurobasal medium (Gibco, Thermo Fisher, Waltham, MA, U.S) with B27 supplement (Gibco, Thermo Fisher, Waltham, MA, U.S). Cell cultures were maintained at 37°C in a 5% CO_2_ humidified atmosphere.

### Culture of human induced pluripotent stem cells (hiPSc)

hiPSc were cultured on a feeder layer of irradiated mouse embryonic fibroblasts (iMEF) in iPSc media (DMEM/F12 (Gibco, Thermo Fisher, Waltham, MA, U.S) supplemented with NEAA (Gibco, Thermo Fisher, Waltham, MA, U.S), Glutamax (Gibco, Thermo Fisher, Waltham, MA, U.S), 10 % KSR (Gibco, Thermo Fisher, Waltham, MA, U.S), 100 μM 2-mercaptoethanol (0.25 mg/ml, Sigma-Aldrich), 10 mM HEPES (0.25 mg/ml, Sigma-Aldrich), 20 ng/ml bFGF (Peprotech, Cranbury, NJ. U.S.) and Pen/Strep (Gibco, Thermo Fisher, Waltham, MA, U.S) and maintained at 37°C in a 5% CO_2_ atmosphere. Media was replaced every two days and cells were splitted every 5 - 6 days based on colony growth. Differentiating colonies were removed from the plate prior to splitting. For passaging, cells were dissociated with collagenase.

### Neuronal differentiation from hiPSc

For the induction of forebrain neurons, hiPSc were differentiated using an embryoid body-based protocol (Zeng et al., 2010). Briefly, hiPSc colonies were dissociated and cultured as aggregates in suspension in iPSc media with 10 μM Y27632 (Tocris, Bristol, UK) and 4 ng/ml bFGF (Peprotech, Cranbury, NJ. U.S.). After 3 days, aggregates were switched to neural induction media (NIM) (DMEM/F12 (Gibco, Thermo Fisher, Waltham, MA, U.S), supplemented with NEAA (Gibco, Thermo Fisher, Waltham, MA, U.S), Glutamax (Gibco, Thermo Fisher, Waltham, MA, U.S), N2 (Gibco, Thermo Fisher, Waltham, MA, U.S), 1 mg/ml Heparin (Sigma Aldrich, San Luis, MO, U.S.), 4ng/ml bFGF and Pen/Strep (Gibco, Thermo Fisher, Waltham, MA, U.S). After 7-10 days, cell aggregates were plated on Geltrex (ThermoFisher, Waltham, MA, U.S.) coated dishes in NIM media with 20 ng/ml bFGF (Peprotech, Cranbury, NJ. U.S.).

Primitive neuroepithelial (NE) structures form over 7-10 days and after 17 days neural rosettes were present. Neural rosettes were selected manually for further expansion on Geltrex coated dishes and grown in neural differentiation media (NDM) (Neurobasal supplemented with NEAA, Glutamax, N2, B27, 1μM cAMP, 10 ng/ml BDNF (Peprotech, Cranbury, NJ. U.S.), 10 ng/ml GDNF (Peprotech, Cranbury, NJ. U.S.), 10 ng/ml IGF-1 (Peprotech, Cranbury, NJ. U.S.), 200 ng/ml ascorbic acid and Pen-Strep. Media was replaced every other day. For final neuronal differentiation, dissociated rossettes were cultured in Geltrex coated coverslips in Neurobasal supplemented with NEAA, Glutamax, N2, B27 and Pen-Strep and were maintained in this media until experimental use. This protocol yields a culture mainly composed of human neurons and astrocytes.

### Cell transfection

N2a and HeLa cells grown at 70-80 % confluence in 35 mm dish were transfected by directly adding to culture media a mixture of 1 μg of indicated DNA and 0.05 mM PEI dissolved in NaCl 0.15 mM, and expression of transfection was performed 24 – 48 h after.

Transfection of human neurons for APP-BACE1 BiFC was carried out in 10 DIV cultures of human neurons derived from hiPSCs growing in 35 mm dish. The conditioned media was replaced by a transfection mixture composed of 8 μg DNA (BACE:CH and APP:VN and BACE1:VC) and 10 μl of Lipofectamine 2000 (ThermoFisher) in 1.0 ml of Optimem (Gibco, Thermo Fisher, Waltham, MA, U.S). After 2 hours, transfection mixture was removed and original conditioned media was restituted. Expression of transfection and BiFC was analyzed after 48 h.

### Immunofluorescence

Immunolabelling was performed as described previously (Bignante et al. 2018). Briefly, cultures grown in coverslips were fixed with 4 %paraformaldehyde, 0.12 M sucrose in PBS for 20 min at 37°C, permeabilized for 5 minutes with 0.2% Triton X-100 in PBS, blocked for 1 hour in 5% horse serum, incubated overnight at 4° C with primary antibody. Afterwards, the coverslips were incubated with Alexa-conjugated secondary antibodies for 1 h at room temperature and 2.5 μM DAPI for 5 min. Finally, coverslips were mounted on slides with Fluorsave Reagent (Calbiochem).

### Analysis of fluorescence and co-localization quantitation

All fluorescent images, including those for co-localization and/or BiFC experiments, were captured in either, an Olympus FV1000 Spectral or an LSM800 Zeiss confocal microscope, equipped with a PLAPON 60X Oil objective. Z-stacks images were acquired with optical cuts of 0.12 μm. Images were deconvolved and projected on Z axis for intensity measurement. Fluorescent intensity was determined using FIJI software (NIH). For colocalization analysis, images were submitted to a background substraction and Manders correlation index M1 was obtained as indicated by Bolte and Cordelières (2006) using the plugin Coloc2 from FIJI software. Manders coefficients were used to calculate the percentage of APP, APPΔβ, APPGP and BACE1 in different organelles. Identical setting for image capture and analysis were conserved across all samples of the same experiment. For primary rat hippocampal neurons, images of cell body and the brightest neurites were acquired and analyzed independently. For human neurons derived from iPSc, images were obtained from the soma and dendrites with a 4x optical zoom, which were identified by MAP2 staining.

For co-localization studies in cell lines, cultures were co-transfected with APP:YFP (or APP mutants) and BACE1:CH and 1 h later cultures were treated as indicated. After 24 h cultures were fixed, immunolabeled against Rab11 or GM130 and fluorescence was analyzed. For analysis of endogenous APP and BACE1 co-localization in primary rat hippocampal neurons, 10-14 DIV hippocampal cultures were treated as indicated and 24 h later were fixed. Endogenous APP, BACE1 and Rab11 were immunolabeled and fluorescence was analyzed.

For fluorescence analysis of BiFC of APP and BACE1 in N2a cells, cultures were co-transfected with APP:VN and BACE1:VC, 1h later were treated as indicated. Depending on the experimental design, cultures were fixed after 24 or 48 h and fluorescence was analysis. For analysis co-localization of BiFC of APP-BACE1 with Rab11 in human neurons, 10 DIV cultures were co-transfected with APP:VN and BACE:VC and 1h later treated as indicated. After 48 h the cultures were fixed and immunolabeled for Rab11 and MAP2 and fluorescence was analyzed.

### Analysis of protein expression by western blot

Cultures of N2a cells were transfected with empty vector (control) or APP:VN and BACE1:VC constructs, and 1 h later the cultures were treated with the γ-secretase inhibitor DAPT (10 μM), gallein (5 μM), o-Aβ (1 μM) or the corresponding vehicles (control). After 48 h, cultures were lysed in RIPA buffer supplemented with protease inhibitors cocktail (SIGMA-FAST, Sigma) at 4° C. Cell lysates were diluted in Laemmli sample buffer, incubated at 95° C for 5 min, resolved in 12 % gel by SDS-PAGE and electrotransferred to nitrocellulose membrane. Thereafter, the membrane was boiled during 5 min in PBS, incubated with agitation overnight at 4°C with anti-APP C-terminal antibody (clone Y188) or anti-α-tubulin (clone B-5-1-2) followed by the corresponding IR800CW/IR680DR secondary antibodies. Thereafter membranes were scanned and visualized using the Odissey system. Quantitative densitometric analysis of the bands was performed using FIJI software. Data from CTFs were normalized with respect to the corresponding from α-tubulin band.

### Characterization of Aβ assemblies by dot blot

2 μl of Aβ monomers (m-Aβ), Aβ fibrils (f-Aβ) or Aβ oligomers (o-Aβ) were seeded onto a nitrocellulose membrane. After of 5 minutes, membrane was blocked with 5% horse serum in PBS for 1 hour and incubated overnight at 4°C with shacking with 4G8, NU1 or NU4 antibodies. Afterwards, membranes were incubated with IR800CW secondary antibody for 1 hour at room temperature and visualized by IR Odissey system.

### Statistical Analysis

Experiments were performed at least in triplicate and independently replicated 2-4 times with similar results. Experimental data were statistically analyzed using t-test or ANOVA followed by LSD Fisher post hoc test and p ≤ 0.05 was consider as statistically significant. Results are presented as Means ± SEM.

## Data availability

The data that support the findings of this study are available from the corresponding author upon reasonable request.

## ACKNOWLEDGEMENTS

The authors thank to Andrea Pellegrini for technical assistance with cell cultures, to Romina Maiorano, Patricio Pereyra, Eliana Martinez, Jesica Piovano, Milagros Nigro and Marisa Gigena for technical assistance with animals from the vivarium, and to Carlos Más and Cecilia Sampedro for technical assistance with microscopes. The authors gratefully acknowledge Dr. Subjohit Roy for BACE1:CH, APP:VN and BACE1:VC plasmids.

This work was supported by grants from ANPCyT PICT 2014-1768 to EAB, ANPCyT PICT2018-02097 and FONARSEC-SB-PBIT 2013-09 to AL; CONICET PIP 11220150100954CO, PUE CONICET 22920160100135CO. EAB and AL are career members of The National Scientific and Technical Research Council (CONICET). MA was supported by fellowship from CONICET.

## Author contribution

MA and EAB designed, carried out and analyzed the bulk of the experiments. PM and AL developed and characterized human cultures. CLF performed colocalization studies and edited the manuscript. GQ helped with microscopy techniques. MFS edited the manuscript. EAB directed the research. EAB and AL conceived the study and wrote the paper.

---

**Supplementary Figure 1.**
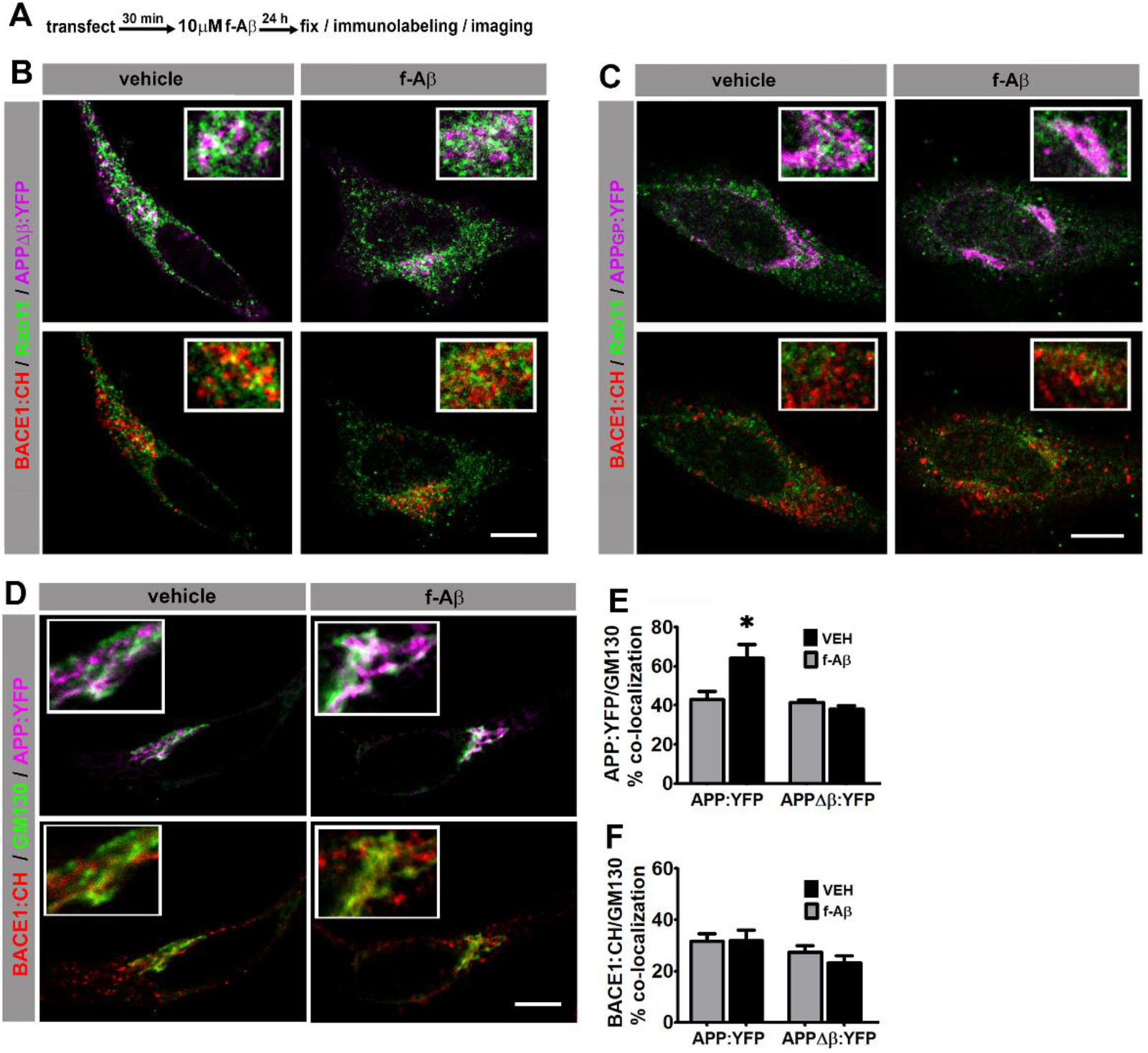
Fibrillar Aβ did not enhance distribution of mutant variants of APP in recycling endosomes and Golgi apparatus. A. Experimental diagram for B-F. Transfected HeLa cells were immunolabeled with antibodies against Rab11 or GM130 followed by AlexaFluor 633-conjugated antibody (Invitrogen) and imaged by confocal microscopy. B and C. Images of HeLa cells co-expressing the indicated proteins and immunolabeled against Rab11. Upper panels show pseudocolor images for Rab11 (green) and APPΔβ:YFP (magenta in B) or APPGP:YFP (magenta in C). Lower panels show pseudocolor images for BACE1:CH (red) and Rab11 (green). Quantitative analysis is shown in Figure 1I and J. D. Images of HeLa cells co-expressing the indicated proteins and immunolabeled against GM130. Upper panel shows pseudocolor images for APP:YFP (magenta) and GM130 (green). Lower panel shows pseudocolor images for BACE1:CH (red) and GM130 (green). E. Manders coefficient of colocalization of APP:YFP and GM130 expressed as percentage. (n = 41; 8-12 for each condition) *: p < 0.001 vs all other conditions by ANOVA followed by Fisher post hoc test. F. Manders coefficient of colocalization of BACE1:CH and GM130 expressed as percentage. (n = 41; 8-12 for each condition) No significant difference between conditions were found by ANOVA. Scale bar 10 μm. Inserts are 2.5 x enlargement of the corresponding image.

**Supplementary Figure 2.**
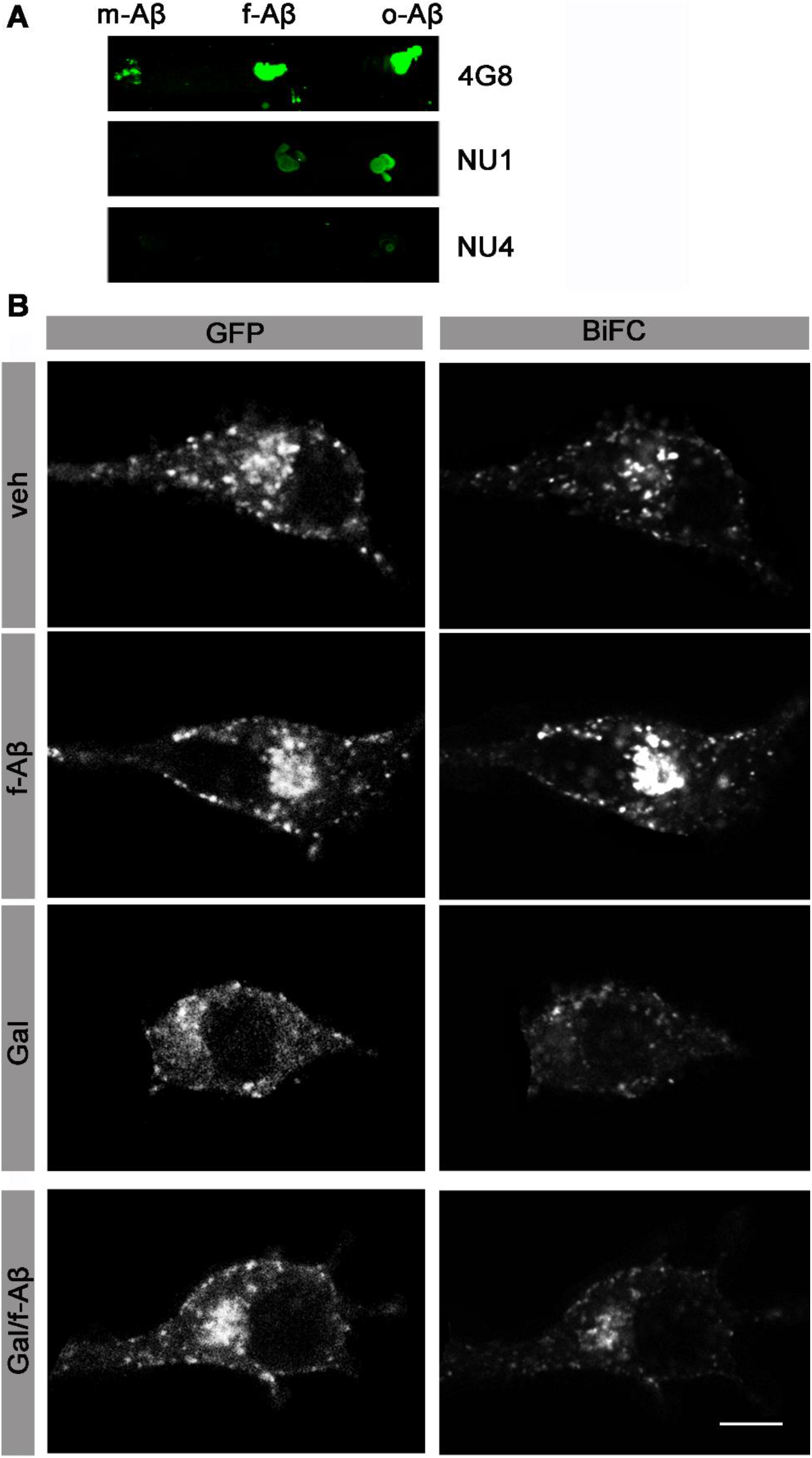
Characterization of Aβ peptide species and control of APP:VN and BACE1:VC expression under treatments. A. Dot blot analysis of Aβ species used in this work. Soluble non-aggregated Aβ 1-42 (m-Aβ), Aβ1-42 oligomers (o-Aβ) and Aβ1-42 fibrils (f-Aβ). Upper panel was revealed with conformation-independent monoclonal antibody 4G8 that recognizes all Aβ species. Middle and lower panels were stained with conformation-dependent antibodies NU1 and NU4, which selectively label oligomeric species. Note that NU1 strongly labeled o-Aβ while weakly recognized f-Aβ. NU4 antibody only labeled o-Aβ. B. Transfected N2a cells with APP:VN and BACE:VC plasmids were treated according left label and immunostained with antibody against GFP (Molecular Probes) followed by AlexaFluor 598 conjugated antibody (Invitrogen) and imaged by confocal microscopy. Left panel depicts GFP stain and right panel BiFC. Note that for similar level of expression of fluorescent protein (anti-GFP stain), level of BiFC intensity varies according treatment. Also, gallein and Aβ treatments did not affect expression of fluorescent protein.

**Supplementary Figure 3.**
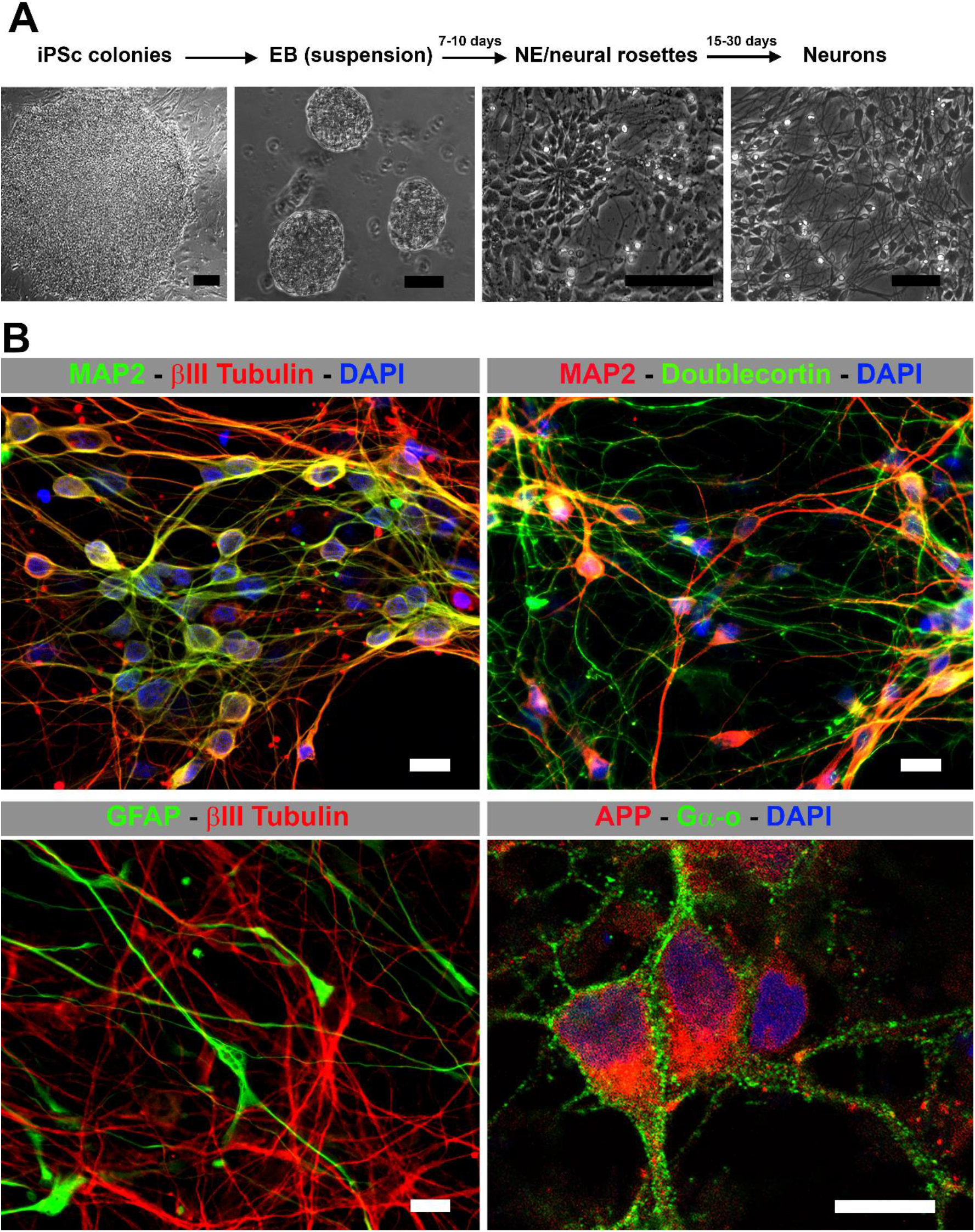
Establishment and characterization of human neuronal cultures. A. Schematic protocol for differentiating human induced pluripotent stem cells (iPSc) to neurons. Shown are phase contrast images of an iPSC colony, embryonic bodies (EB) in suspension, neural epithelium (NE) and neural rosettes and differentiated neurons. Scale bar 50 μm. B. Confocal images of human neuronal cultures at 21 DIV immunolabed for the indicated proteins. Scale bar 10 μm.

